# Paratransgenesis: The dynamics of engineered *Enterobacter* symbionts and Cry1Ac-producing *Enterobacter* for biocontrol of *Helicoverpa* insect pests in crop production

**DOI:** 10.1101/2025.01.01.630992

**Authors:** Abhishek Ojha

## Abstract

The potential application of insect gut microorganisms in paratransgenic crop pest management techniques is being investigated. The *Helicoverpa* gut is known to harbor the symbiont *Enterobacter*. However, little is known about the dynamics of *Enterobacter* in the *Helicoverpa* gut, its interaction with the phyllosphere, and the mechanisms by which it is delivered to this Lepidopteran species. To evaluate the insecticidal activity of Cry1Ac and its capacity to colonize the *Helicoverpa* gut and the tomato phyllosphere, the symbiont *Enterobacter*, which was isolated from the *Helicoverpa* gut, was transformed with constructs 2 (*cry1Ac*-*KmR*-pUC18) and 3 (*KmR*-pGFP). Within 48 hours and 24 hours, respectively, the *Helicoverpa* gut and tomato phyllosphere were effectively colonized by the GFP-producing construct 3-*Enterobacter*. In insect bioassays, Cry1Ac-producing constructs 2-*E. coli* (K12) and 2-*Enterobacter* were used for diet and leaf-dip testing. *Helicoverpa* neonates died 86-100% when fed diets supplemented with 10 -10 cells of Cry1Ac-producing construct 2-*E. coli* (K12), whereas 79–100% died when fed diets supplemented with 10 -10 cells of Cry1Ac-producing construct 2-*Enterobacter*. Furthermore, when the Cry1Ac-producing construct 2-*Enterobacter* (0.10 x 10^9^ cells) was applied to the tomato phyllosphere, it caused 100% mortality of neonates within 192 hours. The successful vector modification, genetic transformation, and establishment of recombinant *Enterobacter* cells in the guts of *Helicoverpa* larvae and on tomato leaves lay the foundation for advancing a paratransgenic strategy. This engineered bacterium could potentially replace synthetic insecticides for managing lepidopteran pests in crops.

## 1. Introduction

A wide variety of insects are known for their adverse impacts on agriculture. Efforts to control these insects are being questioned by the global emergence of insecticide resistance (Robinson et al., 2004). Therefore, it is worthwhile to explore other approaches to complement existing insect control management (Beard et al., 1993). This study demonstrates a paratransgenic strategy introducing foreign molecules into the host insects via symbionts. The plan aims to produce anti-insect gene products in mutualistic (symbiotic) microbes contained by the insects (Chapco and Kelln, 1994; Tang et al., 2004).

Host specificity plays an important role in paratransgenic strategies that help gut microbes transfer foreign (anti-insect) gene products into the insect gut. Native gut microbes are preferred as gene-delivery systems in the paratransgenic method because foreign (non-native) microbes are generally incapable of colonizing a specific gut environment (Chapco and Kelln, 1994; Husseneder and Grace, 2005; Mehta and Murthy, 1992; Thimm et al., 1998; Veivers et al., 1982). Furthermore, genetically engineered microbes firmly connected with a single host insect can reduce potential impacts on non-target organisms. Moreover, the negative impacts frequently depicted when genetically engineered organisms are delivered into competing environmental conditions (Irvin et al., 2004; Van-Elsas, et al., 1994) could potentially be mitigated by employing genes that help the formation of recombinant strains in composite bacterial environments (Hillman, 2002).

In the gut of polyphagous insects like *Helicoverpa armigera* (*H. armigera*) and others, a complex and diverse bacterial population helps in the digestion of plant products (Gayatri et al., 2012; Cazemier et al., 2003; Lemke et al., 2003). The gut bacterial populations that are related to *H. armigera* (Gayatri et al., 2012), *Melolontha melolontha* (Egert et al., 2005), and *Dermolepida albohirtum* (Pittman et al., 2008) are typically observed across the geological allocation of these insect hosts. These commonly revealed gut microbes may play a key role in paratransgenic strategies to control insect pests in crops (Pittman et al., 2008). This could be established by genetically modifying gut microbes to produce anti-insect molecules that adversely affect the insects and employing these microbes in soil infested with insect pests.

Previous studies have revealed the isolation of microbes from the gut of *H. armigera* larvae, as well as from insects collected from different crops and various locations in India, along with an assessment of their diversity (Gayatri et al., 2012). Further, we observed that *Enterobacter* was present in all insect samples (Gayatri et al., 2012). Based on this, we assumed that if *Enterobacter* is capable of colonizing both the insect gut and the leaf surface, it could be a favorable element of a paratransgenic strategy to control insect pests. In this study, we investigate the re-colonization ability of genetically engineered *Enterobacter* (expressing GFP and *KmR*) within the *H. armigera* gut and on the tomato phyllosphere. We also expand this study to include genetically engineered *Enterobacter* (expressing Cry1Ac and *KmR*) to assess Cry1Ac expression within the gut of *H. armigera* and its associated mortality. Furthermore, we examine the lethality of the *Enterobacter* (expressing Cry1Ac and *KmR*) population when applied to tomato phyllosphere, using *H. armigera* neonates in insect bioassays. Our outcomes reveal valuable insights into the potential of using *Enterobacter* as a paratransgenic microbe to control this serious insect pest.

## 2. Materials and methods

### 2.1. Insect larval sampling

The *H. armigera* strain was maintained in the insectary under controlled conditions of temperature (25±5 ^°^C), 70±5% relative humidity (RH), and a 14 h/ 10 h L/D (light/dark) cycle using an artificial diet as described by Sachdev et al. (2014) (Sachdev et al., 2014; Shorey and Hale, 1965). First-instar neonates were then subjected to an insect bioassay. Ten neonates, in triplicates, were used for each insect bioassay experiment.

### 2.2. Recombineering, transformation, and protein expression

A 10 ng *cry1Ac*-pKK223-3 vector (Bacillus Genetic Stock Centre, Columbus, OH) was used as a template for the polymerase chain reaction (PCR, Bio-Rad, USA) to amplify the *cry1Ac* gene. The forward primer 5’-CCCGGGATGTGAACAGTGCCCTTACAA-3’ (which includes an *NdeI* restriction sit) and the reverse primer 5’-CATATGCCACGCTGTCCACGATAAATG-3’ (which includes a *Cfr9I* or *XmaI* recognition site) were employed in the PCR. A PCR was achieved in a 25 µl reaction containing 200 µM dNTPs, 5U of Taq DNA polymerase (New England BioLabs, NEB, Inc., USA), and 200 µM of each primer. The PCR conditions were as follows: 95°C for 2 minutes, followed by 35 cycles of 95°C for 30 seconds, 55°C for 30 seconds, and 72°C for 45 seconds, with a final extension at 72°C for 5 minutes. A 3.0 kb DNA fragment was examined on a 1.5% TAE agarose gel (Sambrook and Russell, 2001) run at 5v/cm. The gel was visualized with ethidium bromide staining, and the DNA was gel-purified using the QIAquick Gel Extraction kit (Qiagen, Inc., Germany) (data not shown) and quantified using a NanoVue Plus spectrometer (data not shown). The purified PCR product was digested with *NdeI* and *Cfr9I* and ligated into the pUC18 vector to achieve construct 1 (*cry1Ac*-pUC18). DH5α competent *Escherichia coli* (*E. coli*) cells were transformed with 10 ng of construct 1. The transformants were selected on agar plates containing ampicillin (100 µg/ml) and subjected to plasmid purification. The colonies were further determined by restriction enzyme digestion and PCR. The *cry1Ac* gene in the recombinant plasmid was sequenced using the Sanger method (data not shown). Further, a 1.5 kb kanamycin-resistance (*KmR*) gene cassette (from pET-28a) was amplified, as described by Chandra et al. (2008) (Chandra et al., 2008; Banai et al., 1985), using the forward primer 5’-CGATCGGATAAACCCAGCGAACCATTTGAG-3’, which includes a *PvuI* recognition site, and the reverse primer 5’-CGATCGCCACGCTGTCCACGATAAATG-3’, which also includes a *PvuI* restriction site. The PCR product was determined on a TAE 1.0% agarose gel (Sambrook and Russell, 2001), and gel-purified using the QIAquick Gel Extraction kit, and quantified (data not shown). The purified PCR product was digested with *PvuI* and cloned into construct 1 and pGFP (ClonTech, Takara Inc.) to achieve construct 2 (*cry1Ac-KmR*-pUC18) and construct 3 (*KmR*-pGFP), respectively. Separate *E. coli* cells were used to transform constructs 2 and 3. The transformants were selected on ampicillin (50 µg/ml) containing agar plates and subjected to plasmid purification. The colonies were then determined by restriction enzyme digestion and PCR. Finally, these constructs (2 and 3) were transformed into both *Enterobacter aerogenes* and *E. coli* K12, isolated from the gut of *H. armigera* (Gayatri et al., 2012). The recombinant construct 3 (*KmR*-pGFP) was expressed by a transformed cell of *Enterobacter aerogenes* and *E. coli* K12 on LA/*KmR* agar and in LB/*KmR* broth, incubated overnight at 37°C.

### 2.3. Protein gel blot analysis

The examination of Cry1Ac from recombinant *Enterobacter aerogenes* (construct 2) and *E. coli* K12 (construct 2) was achieved by applying the Western blot test. The cell pellet was suspended in the extraction buffer (Ojha et al., 2021). The whole-cell lysate was denatured by heating for 10 min at 99°C in 1x SDS-PAGE loading dye. Fifty micrograms (50 µg) of protein from the lysate were separated on a 7.5% SDS-PAGE gel, and the resolved proteins were transferred to a nitrocellulose membrane (0.4µ pore size) (GE Healthcare) by blotting at 180 mA for 45 minutes. The membrane was incubated for 1 h (room temperature, RT) with rabbit anti-Cry1Ac antibody (1:5,000 dilution). It was then incubated for 1 h at RT with goat anti-rabbit IgG-ALP (alkaline phosphatase) antibody (1:3,000 dilution). Finally, color development was achieved using 10 ml of BCIP/NBT (5-bromo-4-chloro-3-indolyl phosphate/nitro blue tetrazolium) substrate solution (Genei Laboratories Pvt Ltd, Bangalore, India). The reaction produced a bluish-grey to black precipitate of Cry1Ac on the membrane.

### 2.4. Feeding of H. armigera larvae with GFP bacteria

To achieve the colonizing potential of *E. aerogenes-*(construct 3) and *E. coli* K12-(construct 3) in the gut of *H. armigera*, a total of 60 first-instar larvae (neonates) were fed a sterile artificial diet supplemented with antibiotics (sterile antibiotic diet) (Brummel et al., 2004) for 48 hrs (2 days). Each larva was fed small cubes (1 g) of the sterile antibiotic diet inoculated with 100 µl/cm^2^ of an overnight-grown bacterial culture of *E. aerogenes-*(construct 3) for 48 hrs. After that, the neonates were transferred to a normal artificial diet. The control group consisted of neonates continuously fed an artificial diet supplemented with *E. aerogenes* (native bacteria, isolated from *H. armigera*). The neonates’ fecal samples were then homogenized in sterile Luria Broth (LB, HiMedia Laboratories Pvt Ltd). A total of 100 μl from the 10^-1^, 10^-2^, 10^-3^, 10^-4^, 10^-5^, and 10^-6^ dilution of GFP-expressing bacteria was applied on LB agar plates containing kanamycin (20 μg/ml). The plates were incubated at 37°C for 24 hrs to examine GFP expression by *E*. *aerogenes-*(construct 3). A similar experiment was performed to examine GFP expression in *E. coli* K12-(construct 3) on LB agar plates containing kanamycin (20 μg/ml). The control group for this experiment consisted of neonates continuously fed an artificial diet supplemented with *E. coli* K12.

### 2.5. Colonization assay with Enterobacter on the tomato phyllosphere

The tomato (*Solanum lycopersicum*) plants used in this investigation were maintained in the greenhouse at the International Center for Genetic Engineering and Biotechnology (ICGEB), New Delhi, India. The tomato leaves (20-25 days old) were cleaned (five times) with sterile water. *E*. *aerogenes*-(construct 3) cells pellets (overnight grown) were washed (thrice) with 20 ml PBS, pH 7.4 (Ojha et al., 2014). The leaves were then sprayed with a suspension of *E*. *aerogenes-*(construct 3) at a concentration of 8 x 10^8^ cells/ ml. The plants were incubated in an enclosed plastic tent on the laboratory bench for 6 hrs at 22°C. Afterward, the plants were transferred to a growth chamber and incubated at 28°C and 95% relative humidity (RH, range 75-99%) with a 12-hour photoperiod for an additional 3 days. Three triplicate leaves from plants were harvested at 24, 96, and 192 hrs. These leaves were washed, five times, with PBS, pH 7.4. The individual leaves were then placed in 15 ml bacterial culture tubes containing 10 ml of LB medium supplemented with 50 µg/ml kanamycin, and incubated for 2 hours at 37°C with shaking at 200 rpm. Fifty microliters of each culture were plated on LB agar plates containing 20 µg/ml kanamycin, and the plates were incubated for 15-17 hrs (overnight) at 37°C. Each experiment was conducted in triplicate and repeated three times.

### 2.6. Diet incorporates bioassays

The diet inclusion method was used to perform the bioassay. *Enterobacter-*(construct 2) was incorporated into the artificial diet at concentrations of 10^5^, 10^6^, 10^7^, 10^8^, and 10^9^ bacterial cells/cm^2^ of diet. Thirty neonates were used for each of the five concentrations of *Enetrobacter-* (construct 2) cells, as well as for the control group, which consisted of neonates on a diet supplemented with 10^9^ *Enetrobacter*-(construct 3) cells/cm^2^ of diet. *Enterobacter* cells were serially diluted in PBS (10 mM, pH 7.4), and (100 µl) of each dilution was added to the semi-synthetic diet in each well of a 12-well plate. The plates were then dried for 1 hour in a laminar airflow workbench. After that, 10 neonates were placed in each well. The lids of all diet plates were perforated with six ventilation holes (2.5 mm diameter). Insect mortality was checked at 24, 48, and 72 hrs. Mortality rates were corrected using Abbott’s formula (Abbott, 1925). In parallel, a Similar experiment was performed with *E. coli* K12-(construct 2) cells at concentrations of 10^5^, 10^6^, 10^7^, 10^8^, and 10^9^ bacterial cells/cm^2^ of diet, with a control group consisting of neonates on a diet supplemented with 10^9^ *E. coli* K12*-*(construct 3) cells/cm^2^ of diet. All bioassay experiments were carried out under controlled environment conditions in the insectary at 25±5 ^°^C, 70±5% relative humidity, with a 14 h:10 h L/D cycle.

### 2.7. Leaf dip bioassay

Laboratory investigations using a leaf dip bioassay approach were executed against the cotton bollworm. Tomato plants (20-25 days old) were used to collect the leaves. The harvested leaves were rinsed twice with sterile water for 10 minutes to eliminate any potential contamination. *Enterobacter, Enterobacter-*(construct 3), and *Enterobacter-*(construct 2) cultures were grown overnight (16-18 hrs, at 37°C, 200 rpm), and the cell pellets were rinsed three times with sterile PBS (pH 7.4). The bacterial cells were then resuspended in 1 ml of PBS, and the optical density was recorded at A_600_ nm. For treatment, 100 µl (0.15 x 10^9^ cells/ml) of *Enterobacter*, 100 µl (0.15 x 10^9^ cells/ml) of *Enterobacter-*(construct 3), and 100 µl (0.10 x 10^9^ cells/ml) of *Enterobacter-*(construct 2) were spread onto the surface of the leaves, which were then dried for 1 h in a laminar airflow workbench. The treated leaves were placed onto 0.8% solidified agarose (10 ml) in Petri dishes. Ten neonates were allowed to feed on each bacterial-treated leaf. This experiment was conducted with four replicates. Insect mortality rates were examined at 24, 48, 72, 96, and 192 hrs. The mortality rates were corrected using Abbott’s formula (Abbott, 1925).

## 3. Results

The green fluorescent protein (GFP)-encoding DNA plasmid (pGFP) revealed good expression in *E*. *coli* K12 and *Enterobacter aerogenes* on ampicillin-containing agar plates. These results also showed that only a small number of untransformed *Enterobacter* cells were present on the ampicillin-containing agar plates, showing that *Enterobacter* cells were resistant to ampicillin (data not shown). Previous reports have demonstrated that *Enterobacter* strains are susceptible to kanamycin (Davin-Regli, 2019). Following this, *Enterobacter aerogenes* cells (carrying the *KmR* gene) were tested on kanamycin-(30 mg/ml)-containing agar plates. No growth of *Enterobacter aerogenes* was observed on the kanamycin-containing agar plates (data not shown), suggesting that *Enterobacter aerogenes* was susceptible to kanamycin.

### 3.1. Colonization of Enterobacter within Helicoverpa gut and on phyllosphere of tomato

A *KmR* gene was introduced into the pGFP plasmid, resulting in a 4689 bp construct, referred to as construct 3 (*KmR*-pGFP) (**Fig. S1**). This construct was then transformed into *Enterobacter*, and pure GFP-expressing cells were obtained (data not shown). This investigation aims to develop a biological expression system for the production of heterologous proteins in *Enterobacter*, utilizing the central transcription enzyme (RNA polymerase, RNAP) and promoters found in natural isolates of the Enterobacterales order. The construct proved useful in colonization studies. The number of transformants of GFP expressing *E*. *aerogenes* was successfully achieved on kanamycin-containing agar plates. Fluorescence from the transformants could be easily visualized using an optical fluorescence microscope and a UV transilluminator. Under normal conditions, all transformants revealed uniform green fluorescence, as observed microscopically and with the UV transilluminator. The fluorescence intensity and optical density scales displayed typical GFP production in *E*. *aerogenes-*(construct 3), with an excitation peak at 488 nm and an emission peak at 513 nm (data not shown).

Neonates were exposed to an artificial diet containing antibiotics for two days. On the third day, the neonates were transferred to an artificial diet containing *Enterobacter-*(construct 3) for two consecutive days. On the fifth day, the neonates were maintained on a diet without *Enterobacter-* (construct 3). Fecal pellets were collected and examined at 24-hrs intervals for the next 7 days to detect GFP-expressing *E*. *aerogenes*. The bacteria were observed to be capable of stable gut colonization for 7 consecutive days (up to the 5^th^-instar stage), suggesting a native colonization characteristic in the gut of *H. armigera* (**Table 1a**). The colonization potential of GFP-expressing *E*. *aerogenes* on the surface of tomato leaves was also determined. GFP-expressing *E*. *aerogenes* was sprayed on the leaves and the number of colonies (50, 110, and 255) was counted at 24, 96, and 192 hrs, respectively (**Table 1b**). These results revealed that *E*. *aerogenes* can effectively colonize both the insect gut and the surface of the leaves.

**Table 1.** Colonization potential of GFP-expressing *Enterobacter* in the gut of insects and on the phyllosphere. (a) Colonization of the intestinal tract of the polyphagous insect *H. armigera* by the GFP-expressing gut microbe *E.* aerogenes. (b) Colonization of tomato leaves by the GFP-expressing gut microbe *E.* aerogenes.

| S. No | Dilution of fecal matter | *Enterobacter* (construct 3) colonies obtained from fecal matter |
| --- | --- | --- |
| 1 | $10^{-1}$ | 95 |
| 2 | $10^{-2}$ | 46 |
| 3 | $10^{-3}$ | 15 |
| 4 | $10^{-4}$ | 2 |
| 5 | $10^{-5}$ | 0 |
| 6 | $10^{-6}$ | 0 |

(b) Colonization of tomato leaves by the GFP-expressing gut microbe *E. aerogenes*.
| S. No | Sampling time (hours) | *Enterobacter* (construct 3) colonies obtained from *the* phyllosphere |
| --- | --- | --- |
| 1 | 24 | 50 |
| 2 | 96 | 110 |
| 3 | 192 | 255 |

### 3.2. Cloning and expression of cry1Ac and kanamycin genes

To achieve construct 2 (*cry1Ac*-*KmR*-pUC18), we amplified 3.0 kb *cry1Ac* and 1.489 kb *KmR* amplicons (data not shown). The 3.0 kb *cry1Ac* gene was cloned into the pUC18 vector (**Fig. 1a**), in-frame with the first methionine of the beta-galactosidase gene in the open reading frame (ORF), resulting in a 5435 bp construct (construct 1: *cry1Ac*-pUC18) (data not shown, **Fig. 1b**).

**Fig. 1.**
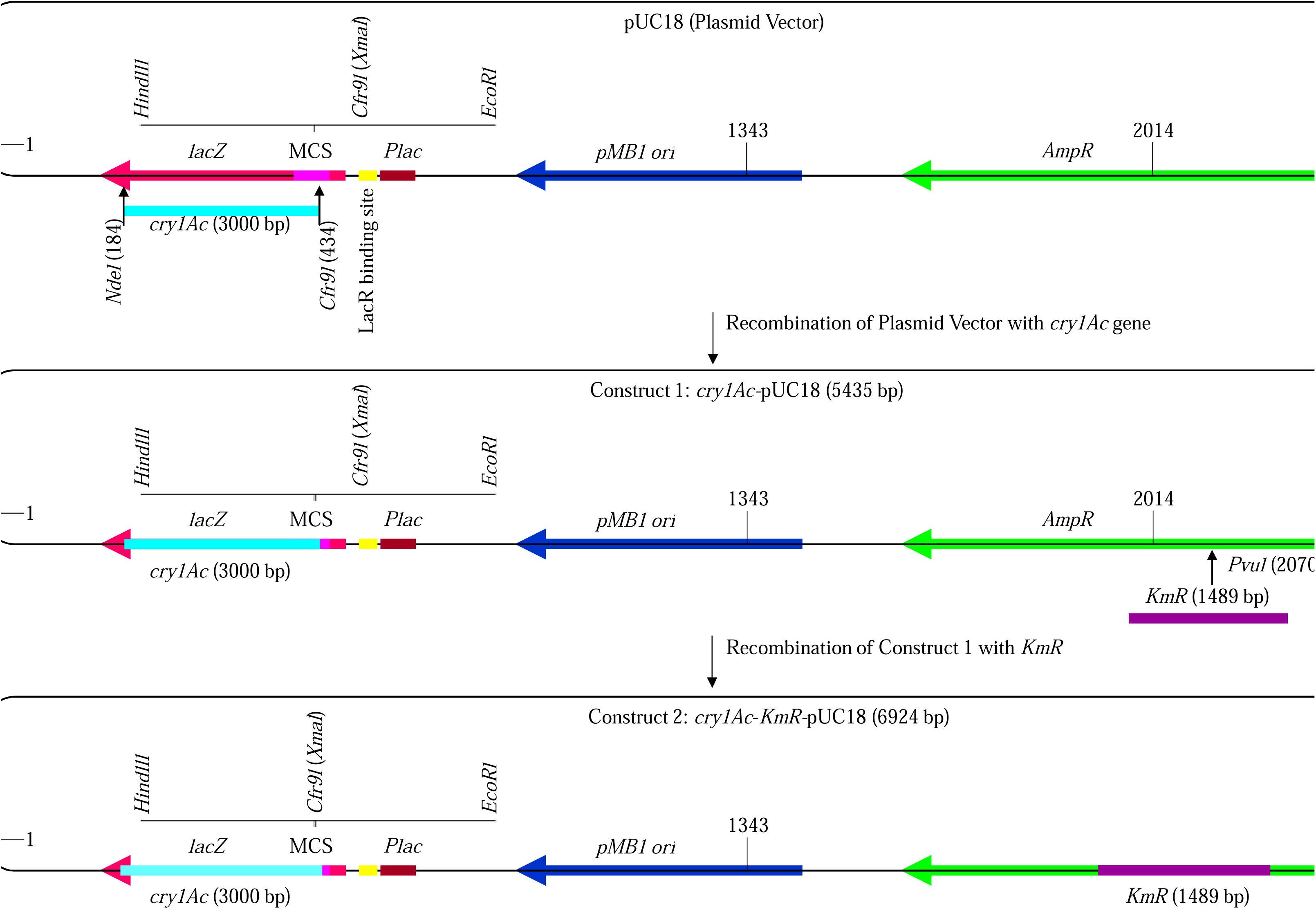
A diagrammatic representation of the structure of the modified pUC18 plasmid vector. The pUC18 vector (a) was modified incorporating the *cry1Ac* (3000 bp, shown in cyan) gene (b) and the *kanamycin* resistance gene (*KmR* 1489 bp, shown in purple), along with the promoter, to produce construct 2 (*cry1Ac*-*KmR*-pUC18) (c).

After then, a 1.489 kb *KmR* DNA fragment was cloned into construct 1 to produce a 6924 bp construct 2 (*cry1Ac*-*KmR*-pUC18) (**Fig. 1c**). The resulting construct 2 was delivered into *Enterobacter* and *E. coli* K12. Cry1Ac expression was revealed in overnight-cultured *Enterobacter*-(construct 2) and *E. coli* K12-(construct 2) cells under the control of the *lac* promoter (**Fig. S2**). The expressed ∼133 kDa Cry1Ac protein was verified by western blotting using anti-Cry1Ac antibody (**Fig. S2**).

### 3.3. Diet and Leaf-Dip Incorporation Insect Bioassay

A bioassay experiment was performed with a total of 420 neonates and overnight-grown bacterial cells [*Enterobacter*-(construct 2), *Enterobacter*-(construct 3), *E. coli* K12-(construct 2), and *E. coli* K12-(construct 3)] (**Table 2**). Five concentrations of Cry1Ac-expressing *E. coli* K12 cells (10^5^, 10^6^, 10^7^, 10^8^, and 10^9^ cells) were individually tested, and all concentrations caused mortality in the neonates. Among the concentrations, 10^6^, 10^7^, 10^8^, and 10^9^ *E. coli* K12 (Cry1Ac-expressing) cells displayed the highest (100%, corrected) mortality, while 10^5^ cells (Cry1Ac-expressing *E. coli* K12 cells) showed the lowest (86%, corrected) mortality (**Table 3**). In contrast, 10^9^ cells of GFP-expressing *Enterobacter* and GFP-expressing *E. coli* K12, which served as negative controls, revealed zero (0%, corrected) mortality (**Table 2a**). Similarly, as a positive control, five concentrations of Cry1Ac-expressing *Enterobacter*-(construct 2) cells (10^5^, 10^6^, 10^7^, 10^8^, and 10^9^ cells) were individually tested, and all concentrations caused mortality. Among these, 10^8^ and 10^9^ *Enterobacter* (Cry1Ac-expressing) cells showed the highest (100%, corrected) mortality, while 10^5^ cells (Cry1Ac-expressing *Enterobacter*) resulted in the lowest (79%, corrected) mortality (**Table 2b**). These outcomes suggest that the gut microbes can be modified to transfer proteins into the hosts.

**Table 2.** Diet incorporates insect bioassays. (a) Cry1Ac-producing *E. coli* K12 (construct 2) and (b) *Enterobacter* (construct 2) were used in the diet incorporating insect bioassay. (a) *E. coli* K12 (construct 3) and (b) *Enterobacter* (construct 3) were used as a control in this bioassay. **Note:** Abbott’s formula was used to calculate the corrected percentage (%) mortality: (= M_observed_ - M_control_ **/** 100 - M_control_) x100. Where *KmR* denotes the *Kanamycin* resistance gene with promoter, and M represents mortality. Construct 2 refers to *cry1Ac*-*KmR*-pUC18, and Construct 3 refers to *KmR*-pGFP.

| Sample name | No. of bacterial cell | Insect used | Mortality | % Mortality | Corrected % mortality |
| --- | --- | --- | --- | --- | --- |
| *E. coli* K12 (construct 3) | $10^9$ | 60 | 3 | 5 | - |
| *E. coli* K12 (construct 2) | $10^5$ | 30 | 26 | 86.66 | 86 |
| „ | $10^6$ | 30 | 30 | 100 | 100 |
| „ | $10^7$ | 30 | 30 | 100 | 100 |
| „ | $10^8$ | 30 | 30 | 100 | 100 |
| „ | $10^9$ | 30 | 30 | 100 | 100 |

(b)
| Sample name | No. of bacterial cell | Insect used | Mortality | % Mortality | Corrected % mortality |
| --- | --- | --- | --- | --- | --- |
| *Enterobacter* (construct 3) | $10^9$ | 60 | 3 | 5 | - |
| *Enterobacter* (construct 2) | $10^5$ | 30 | 24 | 80 | 79 |
| „ | $10^6$ | 30 | 26 | 86.66 | 86 |
| „ | $10^7$ | 30 | 28 | 93.33 | 93 |
| „ | $10^8$ | 30 | 30 | 100 | 100 |
| „ | $10^9$ | 30 | 30 | 100 | 100 |

To examine the potency of *Enterobacter*-(construct 2) in managing *H*. *armigera* populations on tomato leaves, we applied *Enterobacter* (0.15 x 10^9^ cells), *Enterobacter*-(construct 3) (0.15 x 10^9^ cells), and *Enterobacter*-(construct 2) (0.15 x 10^9^ cells) to the surface of detached tomato leaves. A total of 120 neonates (10 neonates per leaf, 4 replicates) were released onto the leaves (**Table 3**). The results showed 100% (corrected) mortality of neonates on *Enterobacter*-(construct 2)-treated leaves, compared to the controls (**Table 3**).

**Table 3.** Leaf dip bioassay. Cry1Ac-producing *Enterobacter* (construct 2) was used to perform the leaf dip bioassay. Wild-type *Enterobacter* and *Enterobacter* (construct 3) were used as controls in this bioassay. **Note:** Abbott’s formula was used to calculate the corrected percentage (%) mortality: (= M_observed_ - M_control_ **/** 100 - M_control_) x100. Where *KmR* denotes the *Kanamycin* resistance gene with promoter, and M represents mortality. Construct 2 refers to *cry1Ac*-*KmR*- pUC18, and Construct 3 refers to *KmR*-pGFP.

| Sample name | No. of bacterial cells | Insects used | Mortality | % mortality | Corrected % mortality |
| --- | --- | --- | --- | --- | --- |
| *Enterobacter* | $0.15 \times 10^9$ | 40 | 0 | 0 | - |
| *Enterobacter* (construct 3) | $0.15 \times 10^9$ | 40 | 0 | 0 | - |
| *Enterobacter* (construct 2) | $0.10 \times 10^9$ | 40 | 40 | 100 | 100 |

## 4. Discussions

Microorganisms are found in almost every environmental niche in nature. The *H*. *armigera* larva gut contains a diverse bacterial population, including both transient species and microbes that are firmly connected with the host. Previous investigations have shown that resident bacteria are ideal candidates for biological gene delivery systems (Pittman et al., 2008; Durvasula et al., 1997; Kuzina et al., 2002). Therefore, this study aimed to deliver recombinant *Enterobacter* expressing Cry1Ac, which can pass through the gut of *Helicoverpa* larvae and withstand the harsh conditions of the gut environment. This study highlights the strong relationship between *Enterobacter* and its host insect, offering the potential for the development of novel insect control approaches in pest management.

We have previously reported (Gayatri et al., 2012) diverse microbes from *H. armigera*. This study establishes *Enterobacter* as a key colonizer of the *H. armigera* gut. This bacterium was found in neonates, regardless of the crop variety or the locality from which they were sampled. In a previous study, *Enterobacter* was also detected in other insect species, further highlighting the significance of its interactions with these pests. This investigation reveals an important relationship between *Enterobacter* and its insect host, providing insights for the development of a promising new strategy for insect control.

These outcomes confirm that, based on the principle of paratransgenesis, the symbiotic bacterium could be a promising tool for controlling *H. armigera*. A previous investigation described the feasibility of using *Enterobacter* from the insect gut to produce toxins (Kuzina et al., 2002). This investigation applied a shuttle vector to genetically modify *Enterobacter* and observed poor production of Cyt1A in the bacterium. Additionally, similar to *B. thuringiensis* and *E. coli,* inclusion bodies (IBs) were detected, although there were notable differences in the IBs expressed in *Enterobacter*.

However, while the *Enterobacter* community could perform as a delivery system, its effectiveness was limited, highlighting that the plasmid DNA designed to transfer the toxin did not function efficiently in *Enterobacter*. Microbes used in such strategies must be easily acquired by the insect during feeding and capable of colonizing the insect gut thereafter (Watanabe et al., 2000). Therefore, it was necessary to investigate the colonization potential of *Enterobacter* in the gut of *Helicoverpa* and on the phyllosphere. *Enterobacter* is similar to *E*. *coli* and both of them are associated with the Enterobacteriaceae family. *E. coli* is currently the best-known experimental organism for genetic manipulation and serves as the workhorse for heterologous protein expression systems (Madigan et al., 2000; Cronan, 2014). Investigations have shown that many of these expression systems are designed for protein production, particularly in engineered *E*. *coli* strains, such as those with T7 promoters or mutations.

The promoters and polymerases mentioned above are not present in native *Enterobacter,* making its application in a paratransgenic strategy unsuitable for producing genetically modified (GM) crops containing *cry* genes, which have become a significant agricultural innovation in modern farming. However, *Bt* cotton crops producing toxins have been established and have confirmed their potential in controlling the insect pest *H*. *armigera* (Abbas, 2018). There are practical challenges in producing GM Cry1Ac or related proteins in a wide variety of crops grown in the field, particularly in developing countries like India. Therefore, the adoption of paratransgenic *Enterobacter* strains could provide a more effective insect control strategy in the field. In this context, our outcomes show that recombinant *Enterobacter* containing the *cry1Ac* gene is capable of sustaining and colonizing healthy leaves, ultimately killing the neonates of *Helicoverpa*.

*Enterobacter aerogenes* isolated from the gut of *H. armigera* larvae (the cotton bollworm), was engineered for use in a paratransgenic strategy. The results revealed that this engineered microbe can effectively colonize within the gut of *H. armigera* larvae and the leaves of tomato plants. Furthermore, the introduction of a modified plasmid carrying the *cry1Ac* gene under the control of the lac promoter enabled the effective production of Cry1Ac *Enterobacter*. When applied to *H. armigera* larvae, this engineered microbe induced mortality. These outcomes highlight the potential of using Cry1Ac as a paratransgenic agent for the effective control of lepidopterous insect pests.

## Declarations

### Ethics approval and consent to participate

Not applicable.

### Consent to publish

Not applicable.

### Availability of data and materials

All data are provided in the main body of the manuscript; materials are available from the authors. All data are included as supplementary files (named as Figure S1 and S2).

## Acknowledgments

The author thanks Raj K. Bhatnagar (International Centre for Genetic Engineering and Biotechnology, ICGEB, India) and Rajagopal Raman (ICGEB, New Delhi, India) for providing access to infrastructure, laboratory assistance, and financial support. Currently, Rajagopal Raman is working at the Department of Zoology, at the University of Delhi, Delhi, India. The author is also thankful to Natarajan Gayatri Priya and Vipin Singh Rana for providing laboratory help for this study at the University of Delhi, Delhi), India.

## Author Information

Insect Resistance Group, International Centre for Genetic Engineering and Biotechnology, Aruna Asaf Ali Marg, New Delhi 110067, India.

## Supplementary materials

**Fig. S1**| Diagrammatic representation of the modification of the plasmid vector pGFP. The pGFP vector was modified by inserting a *kanamycin* resistance gene with a promoter (*KmR*, 1489 bp in purple) to produce construct 3 (*KmR*-pGFP).

**Fig. S2**| Cry1Ac protein immunoblotting analysis. Cry1Ac (∼133 kDa) expression was determined using an anti-Cry1Ac antibody. (1) Molecular weight markers. (2) *E. coli* K12 crude protein (control, without any plasmid DNA), cells collected from the stationary phase of growth (overnight culture). (3) *E. coli* K12 (construct 2) crude protein, cells collected from the log phase of growth. (4) *E. coli* K12 (construct 2) crude protein, cells collected from the stationary phase of growth (overnight culture). (5) *Enterobacter* (construct 2) crude protein, cells collected from the log phase of growth. (6) *Enterobacter* (construct 2) crude protein, cells collected from the stationary phase of growth (overnight culture). (7) *Enterobacter* crude protein (control, without any plasmid DNA), cells collected from the stationary phase of growth (overnight culture). (8) Purified recombinant Cry1Ac protein (positive control). Note: Construct 2 refers to *cry1Ac*-*KmR*-pUC18 for the expression of Cry1Ac protein.

