## Supplementary figures and images for "Paratransgenesis: The dynamics of engineered *Enterobacter* symbionts and Cry1Ac-producing *Enterobacter* for biocontrol of *Helicoverpa* insect pests in crop production"

### Supplementary Figure S1

#### Slide 1
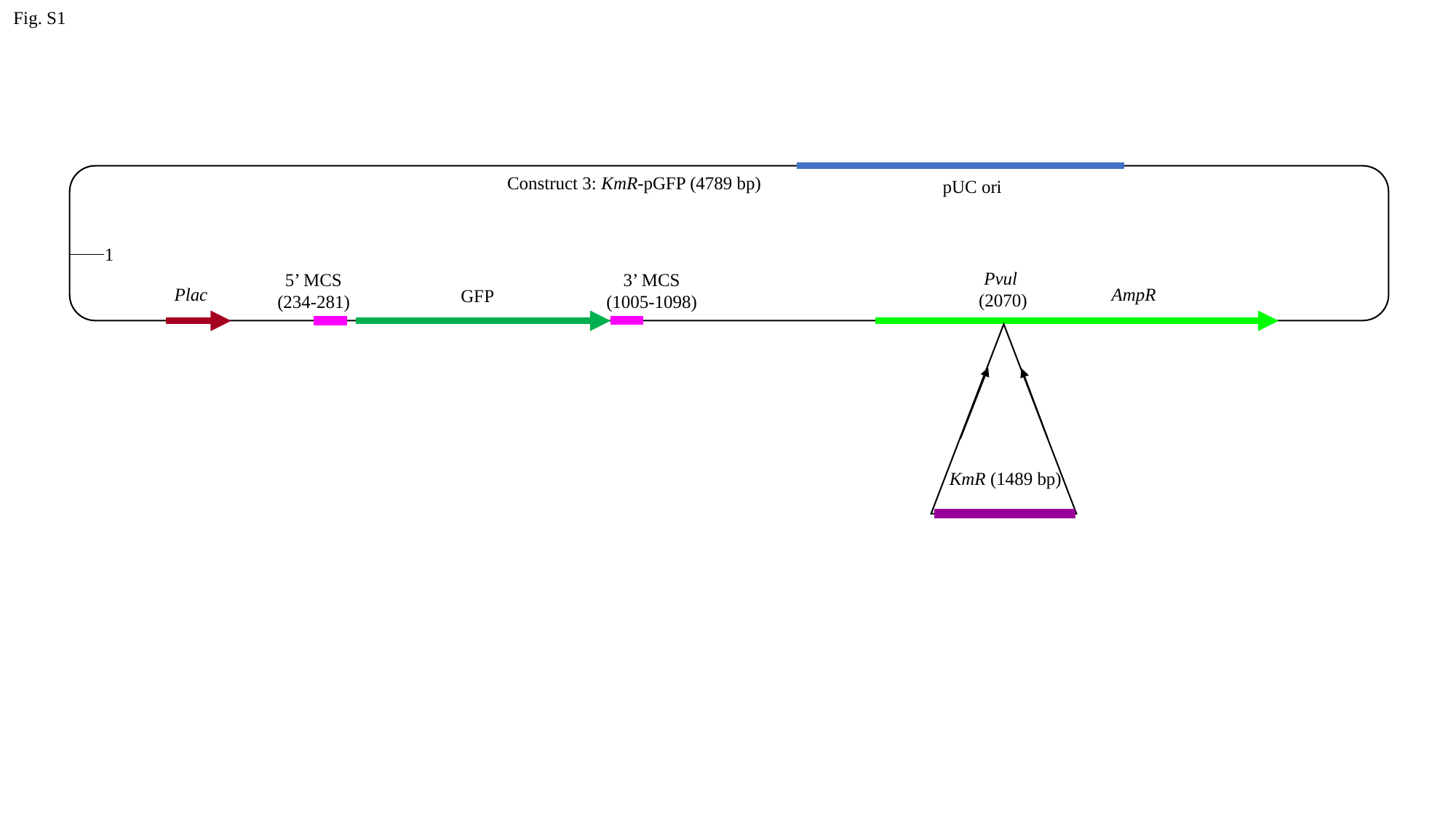

Fig. S1
Construct 3: KmR-pGFP (4789 bp)
pUC ori
1
Pvul
(2070)
5’ MCS
(234-281)
3’ MCS
(1005-1098)
AmpR
Plac
GFP
KmR (1489 bp)
