## Supplementary Figure S2 for "Paratransgenesis: The dynamics of engineered *Enterobacter* symbionts and Cry1Ac-producing *Enterobacter* for biocontrol of *Helicoverpa* insect pests in crop production"

### Slide 1
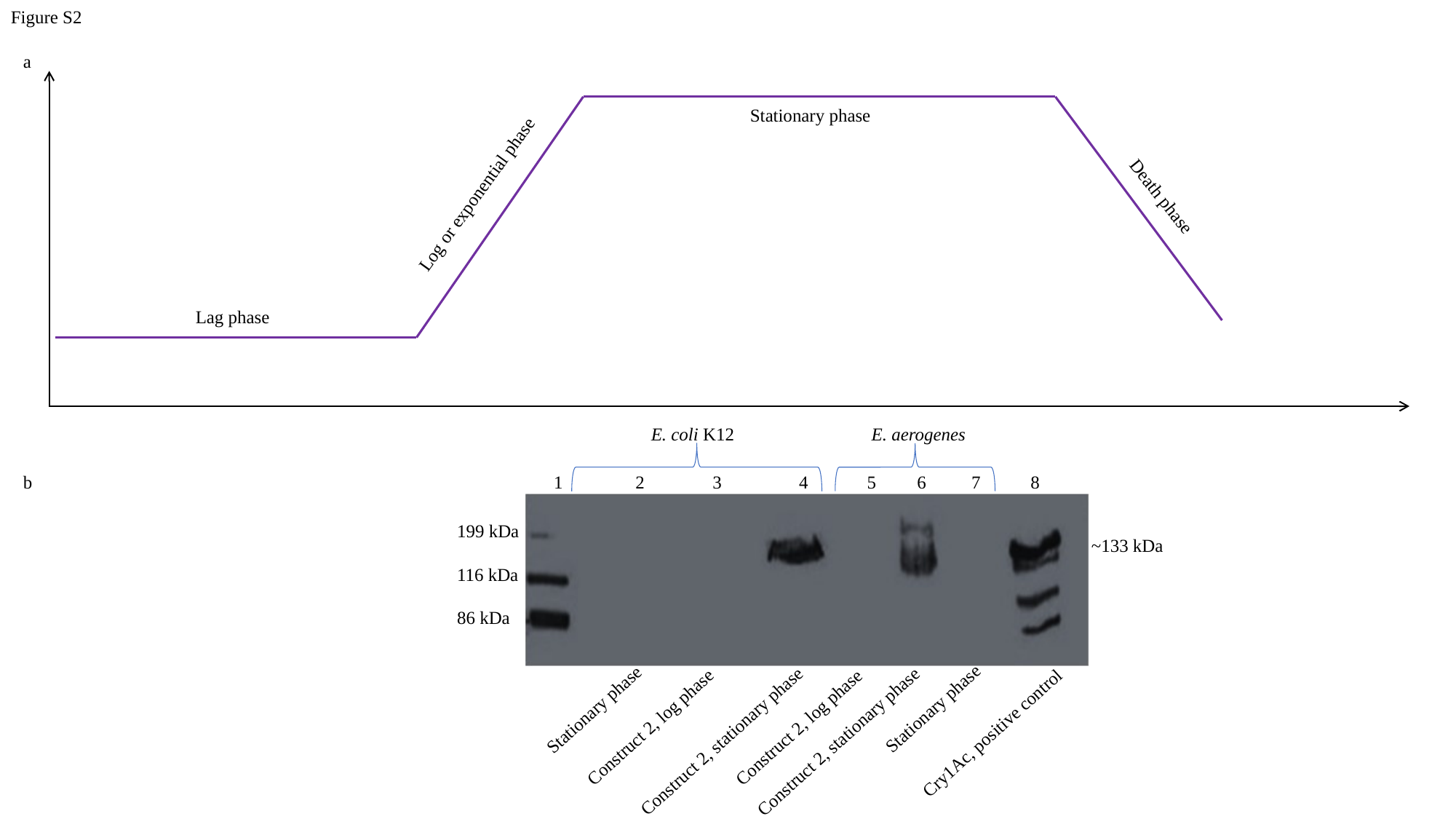

Figure S2
a
Stationary phase
Log or exponential phase
Death phase
Lag phase
E. coli K12
E. aerogenes
1 2 3 4 5 6 7 8
199 kDa
116 kDa
86 kDa
~133 kDa
Stationary phase
Stationary phase
Construct 2, log phase
Construct 2, log phase
Cry1Ac, positive control
Construct 2, stationary phase
Construct 2, stationary phase
b
